# Flexible value coding in the mesolimbic dopamine system depending on internal water and sodium balance

**DOI:** 10.1101/2025.04.16.649233

**Authors:** Takaaki Ozawa, Issei Nakagawa, Yuuki Uchida, Mayuka Abe, Tom Macpherson, Yuichi Yamashita, Takatoshi Hikida

## Abstract

Homeostatic imbalances elicit strong cravings, such as thirst and salt appetite, to restore equilibrium. Although midbrain dopaminergic neurons are known to encode the value of foods, their nutritional state-dependency remains unknown. Here, we show that the activity of the dopaminergic mesolimbic pathway flexibly expresses the positive and negative values of water and salt depending on the internal state in mice. Mice showed behavioral preference and aversion to water and salt depending on their internal water and sodium balance. Fiber photometry recordings revealed that dopamine neurons in the ventral tegmental area and dopamine release in the nucleus accumbens flexibly showed bidirectional excitatory and inhibitory responses to water and salt intake in a state-dependent manner. Furthermore, these dopaminergic and behavioral responses could be simulated by a homeostatic reinforcement learning model. Our results demonstrate the nutritional state-dependency of value coding in mesolimbic dopamine systems, providing new insights into neural circuits underlying homeostatic regulation of appetitive and avoidance behaviors.

## Introduction

Homeostasis refers to the body’s ability to maintain a constant internal environment^1^, and its imbalance can lead to various appetitive behaviors that facilitate its restoration. Typical examples of such cravings include thirst and salt appetite, each of which is triggered by a reduction in the amount of water or sodium concentration in the body, respectively^2–5^. Both water and sodium deficiencies activate the endocrine system, including the renin-angiotensin-aldosterone system (RAAS), which inhibits water and salt excretion and further stimulates appetitive behaviors aimed at restoring balance^5^. On the other hand, it is known that thirst and salt appetite are controlled by distinct neural circuits. Thirst is strongly regulated by the activities of circumventricular organs (CVOs), such as the subfornical organ (SFO) and organum vasculosum of lamina terminalis (OVLT), which are located around the third ventricle and lack a blood-brain barrier^6–9^. The signals from CVOs converge in the hypothalamic median preoptic nucleus (MnPO) to trigger water seeking/taking behavior through a distributed neural network^7,10^. In contrast, salt appetite is controlled by the excitatory activities of specific neural populations in hindbrain and midbrain regions, including the nucleus of the solitary tract (NTS)^11,12^, the pre-locus coeruleus (pre-LC)^12^, and the parabrachial nucleus (PBN)^13^, as well as CVOs^6^.

Dopamine, a key neurotransmitter in the brain’s reward system, is released from dopaminergic neurons centrally located in midbrain regions including the ventral tegmental area (VTA) of the midbrain, which strongly projects to the striatum. These neurons are well-known to play a crucial role in feeding behavior^14^ and are activated by rewarding stimuli, such as food intake under a hunger state^15^. Conversely, they are largely suppressed by aversive stimuli, such as pain^16,17^. This bidirectional response to reward and punishment forms the basis of a prominent theory that dopaminergic activity encodes the value of the stimulus^18^. Moreover, dopamine neuron activity in the midbrain is suggested to represent information analogous to that of reward prediction error (RPE), the difference between the predicted and resulting outcome, in reinforcement learning models^18–20^. Notably, dopamine release in the ventral striatum, including the nucleus accumbens (NAc), in response to reward^21^ and punishment^16^, as well as during the encoding of prediction error-like information^22^, closely mirrors the activity of midbrain dopamine neurons.

Dopamine neuron activity in response to water and salt ingestion is highly dependent on the internal balance of water and sodium in animals. For example, thirst increases the dopaminergic response to water intake^23,24^, while low body sodium levels result in an elevated dopaminergic response to salt intake^23,25,26^. However, it remains unsolved whether excessive water or salt intake suppress dopaminergic activity when the body already contains sufficient water and salt, and whether dopamine neurons in the same animal flexibly show bidirectional responses, such as an excitatory response to rewarding water- or salt-intake and an inhibitory response to aversive water- or salt-intake, in an internal-state-dependent manner. Interestingly, previous experiments have shown that some dopamine neurons in the VTA exhibit excitatory responses to aversive or salient external stimuli, and that dopamine release in specific subregions of the NAc also increases in response to such stimuli ^16–18,27–29^. Moreover, optogenetic activation of midbrain dopamine neurons reduces salt consumption in desalted mice^30^, challenging the idea of purely state-dependent value coding in dopamine neurons.

The objective of this study was to investigate the influence of water and salt imbalance on the activity of the mesolimbic dopaminergic system, and to identify whether bidirectional regulation of dopaminergic responses occurs during water or salt consumption depending on the internal balance. To tackle this, we developed a novel single-drop brief access test to show that mice subjected to water- or salt-restricted conditions exhibit preferences to and avoidances from water or salt depending on their homeostatic needs. Firstly, we demonstrated that our behavioral task was suitable for measuring hedonic and aversive behavioral responses to sweet and bitter outcome, respectively, as well as for quantifying excitatory and inhibitory responses of dopamine neuron in the VTA. Next, through recording of dopamine neuron activity in the VTA and dopamine release in the NAc using fiber photometry, we found that mesolimbic dopaminergic responses were bidirectionally regulated in response to water or salt intake depending on internal water or salt levels. Furthermore, we found that this state-dependent and bidirectional activity of dopaminergic circuits could be qualitatively simulated by a Homeostatic Reinforcement Learning (HRL) model^31,32^. Our results strongly suggest that dopaminergic activity reflects the value of food in relation to the water and mineral balance in the body.

## RESULTS

### Behavioral and dopaminergic responses to appetitive and aversive outcomes

In this study, we first developed a single-drop brief access test to measure liquid food palatability and dopaminergic reactivity in mice. Specifically, we investigated licking responses to sucrose (sweet) and quinine (bitter) solutions in water-restricted mice. During the test, we recorded the activity of midbrain dopamine neurons in response to liquid consumption in freely-moving animals. This was achieved by virally expressing the fluorescent calcium sensor jGCaMP8s in a cre-dependent manner in VTA dopamine neurons of DAT-cre mice, and observing fluorescence through an implanted optical fiber (Fig. 1ab). In the single-drop brief access test, mice could earn either one drop (10ul) of sucrose (300mM) or quinine (1mM) by licking a drinking spout (Fig. 1cd).

**Figure 1.**
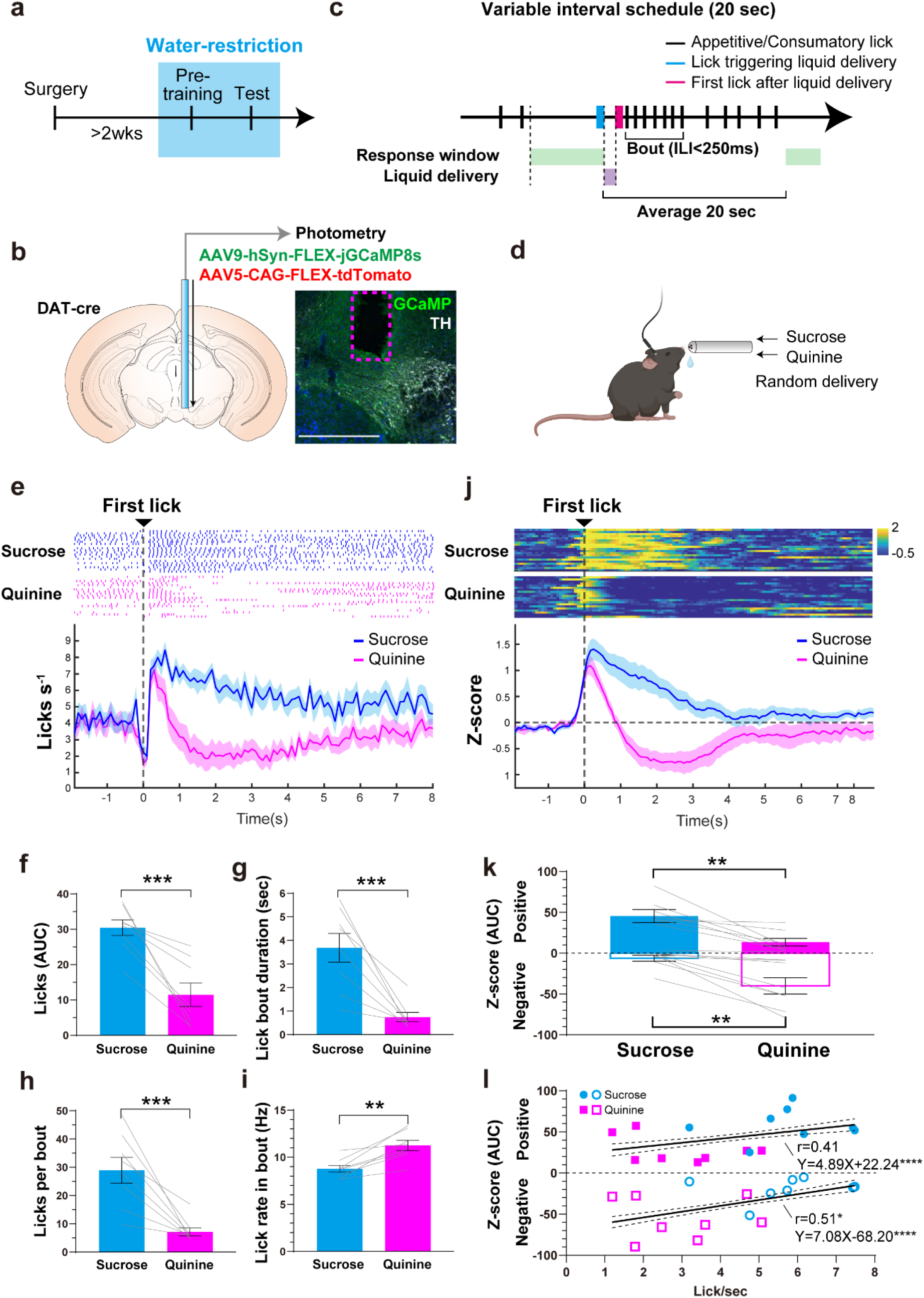
Licking responses and neural responses of midbrain dopamine neurons to sweet and bitter tastants. a Schedule of training and testing in a single-drop brief access test. b Fiber photometry recording of calcium activity in VTA dopamine neurons. c Structure of our single-drop brief access test using a VI20 schedule. The first lick in the response window was followed by liquid (sucrose or quinine) delivery. d In-vivo fiber photometry recording of neural responses in dopamine neurons evoked by liquid ingestion in free-moving mice. e Peri-event time histograms (PETHs) of averaged licking behavior and the raster plots of licking exhibited by an example animal. f Analysis of area-under-the-curve (AUC) of consummatory licks. g Lick bout duration (sec). h Licks per bout. i Lick rate in bout (Hz). j PETHs of the averaged z-score of dopamine neuron activity. Heatmaps show the activity of dopamine neurons in an example animal. k AUCs of dopamine neuron activity evoked by liquid ingestion. l Correlation and liner regression analysis between licking and dopamine responses. Gray lines overlayed on bar plots indicate data from individual animals. **** p<.0001, *** p<.001, ** p<.01, and *p<.05. l Asterisks beside regression equations show significant differences of the slope from zero.

Analysis of licking behavior revealed that mice showed preference or aversion to sucrose or quinine, respectively, in the single-drop brief access test. In peri-event-time-histograms (PETHs) of licking responses during liquid consumption (Fig.1e), it was revealed that mice showed sustained licking responses to sucrose but immediate cessation of licking when quinine was delivered (Fig.1e). Area-under-the-curve (AUC) analyses of licking responses (Fig.1f) demonstrated that mice showed greater licking responses to sucrose than to quinine. Furthermore, the analysis of the licking microstructure ^33,34^ revealed that mice showed longer bout durations (Fig.1g), a larger number of licks per bout (Fig.1h), and a smaller lick rate in each bout (Fig.1i) during sucrose consumption compared to those of quinine. Collectively, these results suggest that sucrose has a higher palatability than quinine, and that the palatability of a liquid can be accurately measured using our task.

Fiber photometry recording revealed a bidirectional change in VTA dopamine neuron activity in response to sucrose and quinine (Fig.1jk, S1, S2). PETH analysis revealed that the activity of VTA dopamine neurons was increased in response to sucrose intake, and transiently increased but immediately reduced to below zero in response to the ingestion of quinine (Fig.1j). This multiphasic response, an initial excitation followed by suppression, of midbrain dopamine neurons to aversive stimuli is in line with that reported in a previous electrophysiology study^35^. Due to this multiphasic response pattern of dopamine neurons after liquid consumption, we separately analyzed the positive and negative components of dopamine neuron activity AUCs^36^ (Fig.1k). We found that the positive component of dopamine neuron activity change during sucrose intake was significantly higher than that of salt intake, whereas the negative component was larger during quinine consumption. We also analyzed the covariance between licking responses and dopamine neuron activity change by correlation and liner-regression (Fig.1l), and found that licking was positively correlated with dopamine neuron activity. Taken together, these findings indicate that our behavioral task in combination with fiber photometry in-vivo imaging was suitable to assess the palatability of liquid stimuli and the resulting dopaminergic response.

### State-dependent, bidirectional responses of dopamine neurons in the VTA to water and salt

Next, we tested state-dependent preference or aversion to water or salt in the same animal by manipulating the internal water or sodium level in mice through dehydration or salt-restriction (Fig. 2abc, S3). As with the previous experiment, DAT-cre mice were infused with a cre-dependent jGCaMP8s into the VTA, above which an optic fiber was implanted for recording of fluorescence (neural activity) (Fig. 2b). Mice then underwent a period of water-restriction (water-restriction phase 1), followed by a period of salt-restriction (salt-restriction phase), followed by a second period of water-restriction (water-restriction phase 2) to evaluate the effect of different internal states on motivated behavior and dopamine neuron activity (Fig. 2a). During these water- or salt-restricted states, mice could earn either one drop (10ul) of fresh water (water) or salt-water (salt; 300mM or 750mM) by licking the drinking spout (Fig. 2c).

**Figure 2.**
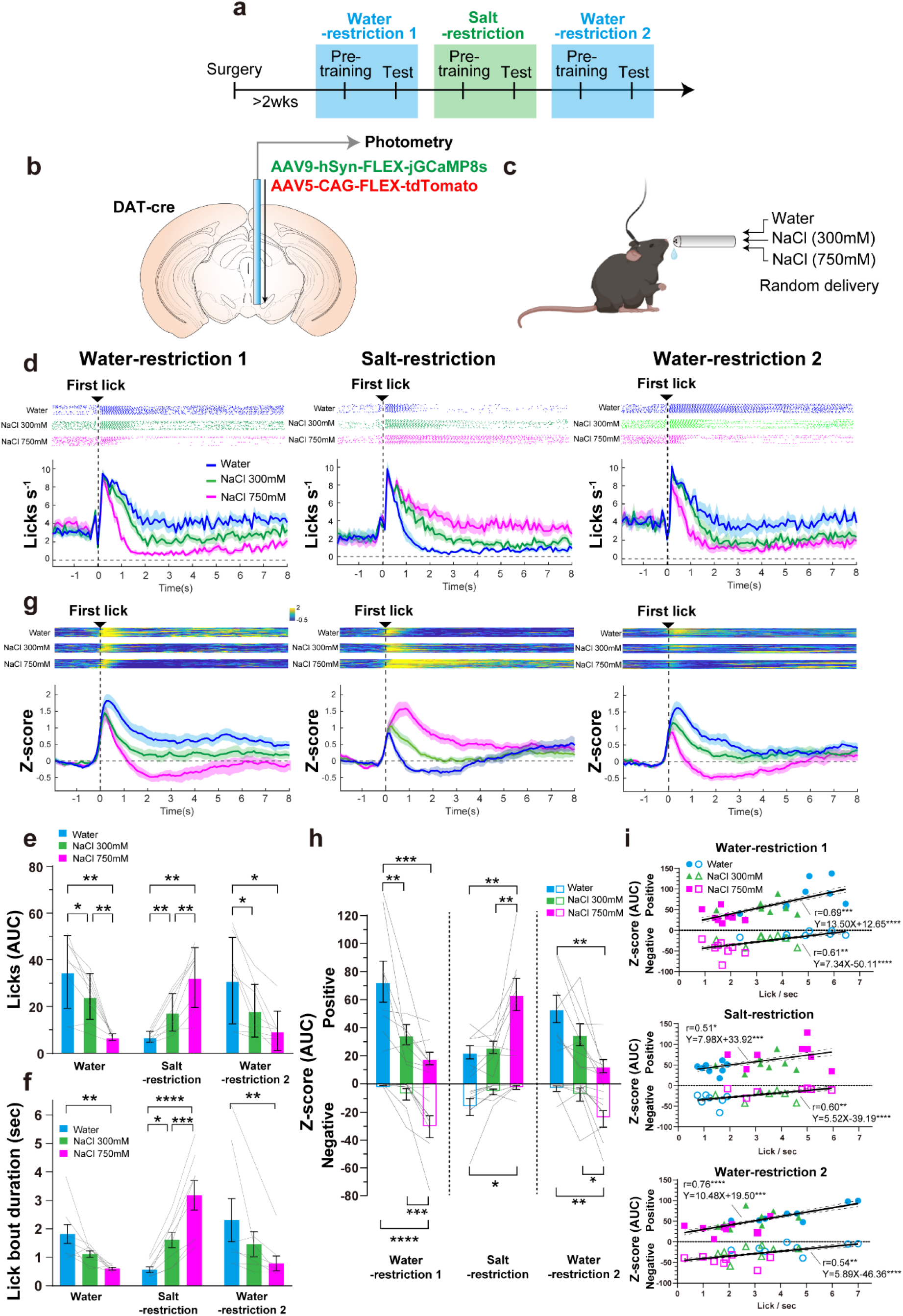
State-dependent preference/aversion and bidirectional responses of midbrain dopamine neurons to water or salt intake. a Schedule of training and testing for state-dependent preference and aversion to water or salt. b Fiber photometry recording of calcium activity in VTA dopamine neurons. c In-vivo fiber photometry recording of neural responses in dopamine neurons evoked by liquid ingestion in free-moving mice. d PETHs of averaged licking behavior and raster plots of licking exhibited by an example animal. e AUCs of licking responses. f Lick bout duration (sec). g PETHs of the averaged z-score of dopamine neuron activity with the heatmaps of an example animal. h AUCs of dopamine neuron activity evoked by liquid ingestion. i Correlation and liner regression analysis between licking and dopamine responses in each phase. Gray lines overlayed on bar plots indicate data from individual animals. f, g, h post-hoc Tukey’s multiple comparisons test. **** p<.0001, *** p<.001, ** p<.01, and *p<.05. i Asterisks beside regression equations show significant difference of the slope from zero.

Analysis of licking behavior revealed that mice showed preference or aversion to water or salt depending on the water or salt restricted state (Fig.2d, f, g). In water-restriction phase 1, mice showed sustained licking responses to water (Fig.2d, left). Conversely, they started but immediately stopped licking when salt was delivered (Fig.2d, left). In the salt-restriction phase, mice showed more licking in response to salt delivery than to water delivery (Fig.2d, middle). In water-restriction phase 2, we again observed sustained licking to water delivery and immediate cessation of licking to salt delivery (Fig.2d, right). Next, we analyzed the AUCs of licking behavior (Fig.2e) and the licking microstructure (Fig.2f, S4). This analysis revealed that mice showed more licking responses to water compared to salt under water-restriction and more licking responses to salt when desalted. Furthermore, we also confirmed that increased licking responses to salt under salt-restriction disappeared when the epithelial sodium channel (ENaC) was blocked by amiloride^12^ (Fig.S5).

Fiber photometry recording revealed a state-dependent and bidirectional change in VTA dopamine neuron activity in response to water or salt, suggesting flexible value coding based on homeostatic needs (Fig.2g, h, i, S6). In water-restriction phase 1, the activity of dopamine neurons was increased in response to water intake and was transiently increased but immediately reduced to below zero in response to the ingestion of high salt (Fig.2g, left). Interestingly, this relationship between liquid intake and dopamine neuron activity was reversed in the salt-restriction phase, where activity increased during salt intake but was inhibited during water intake (Fig.2g, middle). Finally, in water-restriction phase 2, we observed an increase in dopamine neuron activity during water consumption and a transient increase followed by strong inhibition of activity during salt consumption, similar to that observed during water-restriction 1 (Fig.2g, right). Analysis of the AUCs of dopamine neuron responses revealed that, under both water-restriction phases, the positive component of responses to water intake was significantly higher than that to salt intake. Conversely, under salt-restricted conditions, dopamine neuron responses were higher during salt intake than during water intake (Fig.2h). Notably, the negative component of dopamine neuron activity during salt ingestion was larger than that during water under both water-restricted conditions, and was larger during water consumption than that during salt consumption when animals were salt-restricted (Fig.2h). Importantly, we also found a significant positive correlation between licking and dopamine neuron responses (Fig.2j) in each phase suggesting that VTA dopamine neurons encode state-dependent value signals linked to consummatory behavior.

### State-dependent, bidirectional dopamine release in the nucleus accumbens in response to water and salt

Next, we measured changes of dopamine release levels in the NAc core^25^ by taking advantage of the dopamine sensor, GRAB-DA2m (Fig. 3ab, S7). Analysis of licking behavior confirmed the replication of state-dependent preference and aversion to water and salt in mice, consistent with findings from the previous experiment. PETH analysis revealed that mice showed sustained licking behavior in response to the consumption of water while their licking ceased immediately when salt was delivered under water-restriction phases (Fig.3c, left, right). Conversely, under salt-restriction, mice showed higher licking responses to salt compared to water (Fig.3c, middle). Analysis of AUCs and the licking microstructure revealed that mice showed greater licking responses to water compared to salt under water-restriction, and more licking responses to salt under salt-restriction (Fig.3d, e, S8).

**Figure 3.**
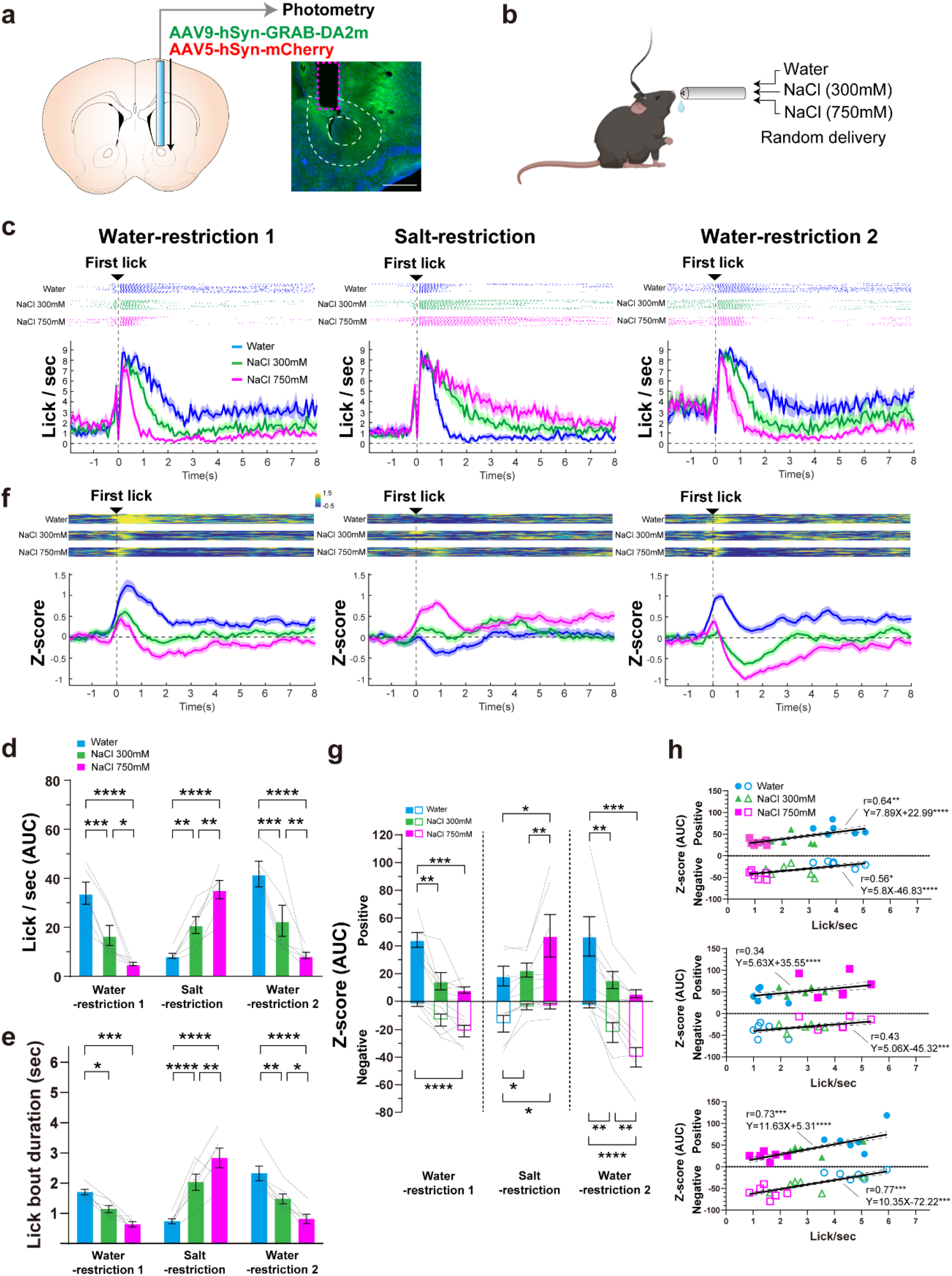
State-dependent bidirectional change of dopaminergic responses to water or salt ingestion in the NAc. a Fiber photometry recording of dopamine changes in the NAc. b In-vivo fiber photometry recording in free-moving mice. c PETHs of averaged licking behavior and raster plots of licking exhibited by an example animal. d AUCs of licking responses. e Lick bout duration (sec). f PETHs of the averaged z-score of the dopamine change with the heatmaps of an example animal. g AUCs of dopamine changes in response to liquid ingestion. h Correlation and liner regression analysis between licking and dopamine responses in each phase. Gray lines overlayed on bar plots indicate data from individual animals. d, e, g post-hoc Tukey’s multiple comparisons test. **** p<.0001, *** p<.001, ** p<.01, and *p<.05. h Asterisks beside regression equations show significant difference of the slope from zero.

Fiber photometry recording revealed nutritional state-dependent, bidirectional changes in NAc dopamine in response to water or salt intake, suggesting state-dependent value coding similar to that observed in VTA dopamine neurons (Fig.3f, g, S9). Under water-restriction, dopamine increased in response to water intake but was transiently increased then immediately reduced to below baseline during high-concentration salt (750mM) consumption (Fig.3f, left, right). However, when desalted, dopamine increased in response to salt intake but was reduced to below baseline during water intake (Fig.3f, middle). Analysis of AUCs revealed that under both water-restriction conditions, the positive component of dopamine was significantly higher during water intake than salt intake, whereas under salt-restriction, it was higher during salt intake than water intake (Fig.3g). In contrast, under both water-restricted conditions, the negative component of dopamine change was larger during salt intake than water intake, while under salt-restriction, the it was larger during water intake than salt intake (Fig.3g). We also assessed the covariance between licking responses and dopamine change using correlation and liner-regression analyses (Fig.3h). As a result, we found that increased licking was positively associated with higher dopamine levels in each water/salt deprivation condition, suggesting a significant relationship between state-dependent value coding by NAc dopamine and consummatory behavior.

### A homeostatic learning model explains state-dependent activity in the dopaminergic system

To better understand the computational principles underlying the dopaminergic responses observed in our animal experiments, we utilized a homeostatic reinforcement learning (HRL) model. This framework enabled us to simulate how internal physiological states, such as water or sodium deprivation, influence value-based decision-making and dopaminergic prediction error signaling. By assessing whether the model could qualitatively reproduce both the behavioral and neural outcomes across deprivation conditions, we aimed to determine whether homeostatic control mechanisms could explain the observed patterns of intake and NAc dopamine activity.

We conducted simulations of the animal experiments under the six deprivation/testing conditions described above and examined the outcomes during water or sodium intake tests (Fig.4ab). Notably, under each condition, the HRL model’s temporal difference (TD) errors (Fig.4c) and estimated number of licks (Fig.4d) qualitatively paralleled dopaminergic activity (Fig.2h, 3g) and licking responses (Fig.2e, 3d), respectively, seen in our animal experiments. This correspondence suggests that the HRL framework can capture key trends in dopaminergic signaling associated with changes in internal states and consummatory behavior.

We describe the model’s behavior in detail for each condition as follows. Under the water-deprived condition tested with water (WD-W), the model successfully reproduced the observed behavioral patterns (Fig.4e). During training, water intake reduced the internal drive (D) as the hydration state moved towards its ideal setpoint, yielding positive reward (R). This increased the state-action value (Q), reinforcing the intake behavior. At the start of the test phase, the model retained this Q-value despite being re-initialized to a deprived state, resulting in positive TD errors upon water consumption. On the contrary, in the WD-300 condition (water-deprived, tested with 300 mM NaCl), while training similarly reinforced water intake (Fig.4f), during the test phase, the beneficial effect of partial hydration was counteracted by the aversive effect of consuming excessive salt, resulting in smaller rewards and smaller positive TD errors relative to WD-W. Finally, in the WD-750 condition (water-deprived, tested with 750 mM NaCl), the high salt content had a stronger negative impact, leading to negative rewards and negative TD errors (Fig.4g).

In contrast to the water-deprived condition, a different pattern of behavior emerged in the model under sodium-deprived conditions. When mice were tested with water (SD-W), it further diluted sodium levels, producing negative rewards and negative TD errors (Fig.4h). In the SD-300 condition (sodium-deprived, tested with 300 mM NaCl), the positive effect of sodium replenishment partially offset the negative effects of excess water intake, resulting in modest positive rewards and TD errors (Fig. 4i). Finally, in the SD-750 condition (sodium-deprived, tested with 750 mM NaCl), the strongly positive effect of sodium replenishment outweighed the negative effects of excess water intake, resulting in high rewards and large positive TD errors (Fig.4j).

**Figure 4.**
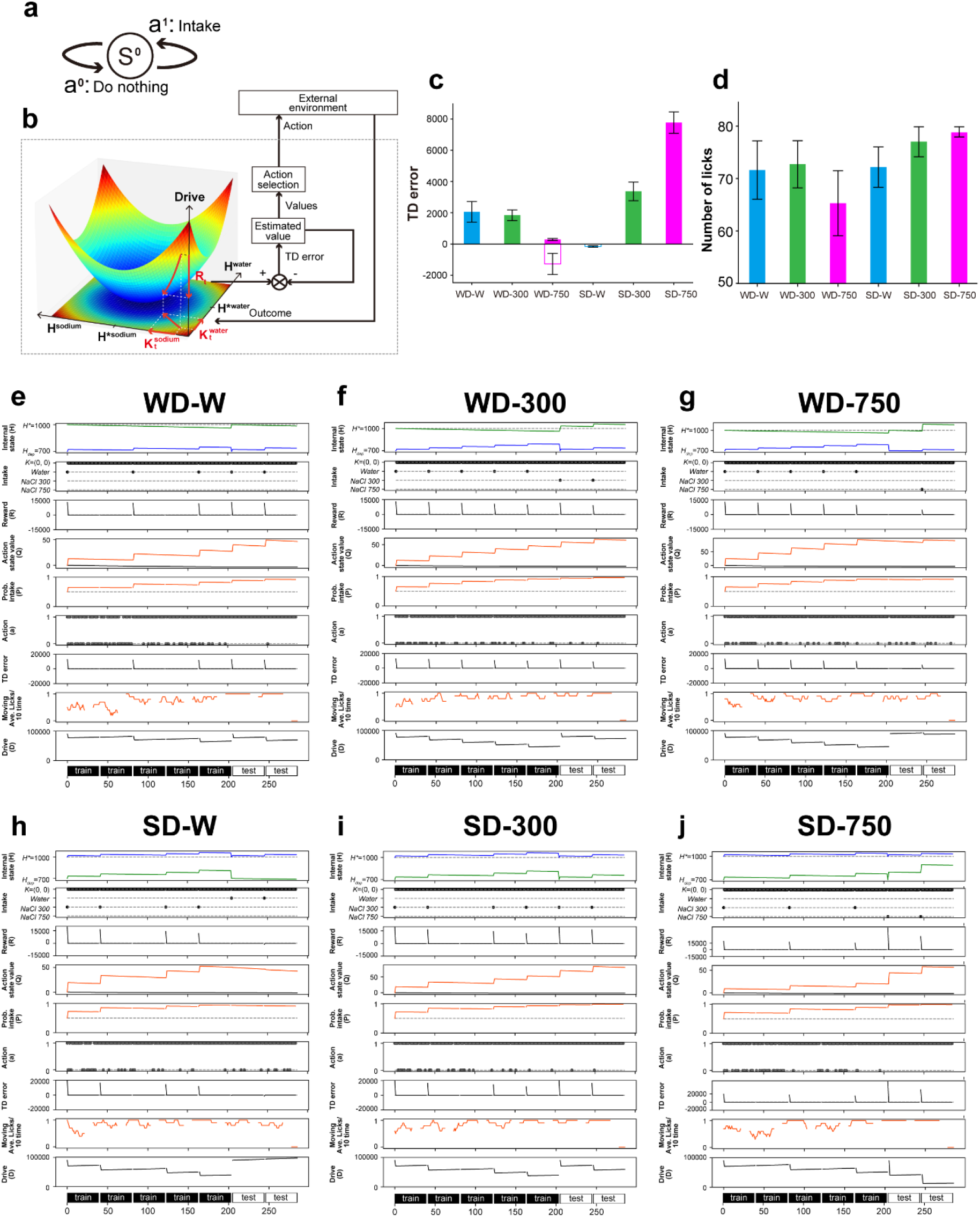
Simulations of state-dependent changes in dopamine and licking responses using the homeostatic reinforcement learning model. a In all simulations, a single (external) state and two actions were defined. b Graphical overview of the homeostatic space and algorithm. c Total temporal difference (TD) error from the tests. d Cumulative licking data from the tests. e-j Time series data for all conditions. The results from five training phases and two test phases are included in a single time series. e Water intake under water deprivation (WD-W), f 300mM salt intake under water deprivation (WD-300), g 750mM salt intake under water deprivation (WD-750), h Water intake under salt deprivation (SD-W), i 300mM salt intake under salt deprivation (SD-300), and j 750mM salt intake under salt deprivation (SD-750) are illustrated. The dashed line for the internal state (H) indicates the setpoint, while the dashed line for the probability of intake (P) represents the threshold at P(Intake) = 0.5. Dotted lines indicate possible discrete variables in Intake and Action (a), serve as reference lines for P = 0, 0.5, and 1 in Probability of intake, and indicate when the value is 0 in Q, TD error, Moving average of lick, and Drive.

Together, these simulation results demonstrate that the HRL model qualitatively captures the dopaminergic dynamics observed in our animal experiments and offers a computational framework for understanding state-dependent value coding.

## DISCUSSION

Although value coding in the mesolimbic dopamine pathway has been reported in previous studies, its nutritional-state-dependency has not been fully substantiated by experimental evidence. In the current study, we developed a single-drop brief access test and successfully revealed both excitatory and inhibitory dopaminergic activity in response to water or salt ingestion depending on the homeostatic needs of the animal. Furthermore, we found that this bidirectional modulation of dopamine signaling could be recapitulated by a HRL model. Together, our experimental and computational results strongly support the idea that value coding in the mesolimbic dopamine pathway is critically shaped by the internal nutritional state.

Previous studies have reported the representation of value information in dopamine neuron activity. Specifically, the majority of midbrain dopamine neurons are activated in response to reward but inhibited in response to punishment^18,20^. It has been found that the increase in dopaminergic response to reward is dependent on the internal state of the animal. For example, water intake during dehydration^23,37^, food intake during fasting^15,21^, and salt intake under sodium restriction^25,38–40^ all induce excitatory responses in dopamine neurons, indicating that consumption to satisfy the homeostatic needs acts as a powerful reward. In the present study, mice exhibited strong preferences for water during water restriction and for salt intake during salt restriction. Furthermore, we also found that strong activation of VTA dopamine neurons and dopamine release in the NAc were elicited during these reward intakes. These results support the state-dependent nature of reward-induced dopamine neuron activation that was reported in previous studies. However, it has also been suggested that distinct populations of dopamine neurons can encode the saliency of stimuli (i.e. the absolute magnitude of value), and that these neurons are activated not only in response to rewarding but also to novel or aversive stimuli^17,18,27,28^. Therefore, it is necessary to investigate neural responses to both reward and punishment in order to examine the representation of value in dopamine neurons. In the present study, we demonstrated that salt intake during water-restriction and water intake under salt-restriction suppressed consumption behavior in mice, and that intake of these aversive stimuli caused inhibition in VTA dopamine neurons and reduced dopamine release in the NAc. Our results suggest that both reward-induced excitatory responses and punishment-induced inhibitory responses in the mesolimbic dopamine pathway are strongly dependent on the water and salt balance in the body. Overall, our results provide the first direct evidence that dopaminergic encoding of value is dependent on internal physiological state.

The specific neural circuits that drive the excitatory dopaminergic responses to the ingestion of water or salt reward, depending on the homeostatic needs, remain unidentified. According to previous studies, water ingestion during the dehydration has been shown to increase dopaminergic activity through both oral sensation ^41^ and gastrointestinal signaling ^42^. It was reported that the immediate oral sensation of water triggers a rapid activation of dopamine neurons, while the slower absorption of water in the digestive system causes a gradual increase in midbrain dopamine activity, which can take several minutes to occur ^42^. Given that the dopaminergic response observed in our study emerged within a few seconds, it is plausible that this excitatory dopaminergic response was driven by rapid sensory signals triggered by the detection of water in the oral cavity and/or the throat, and then transmitted through the trigeminal nerve to the central nervous system ^43^. It has been shown that activity in lamina terminalis (LT) regions, (SFO, MnPO, and OVLT) cause thirst, and that the activation of excitatory neurons in the SFO is especially critical for dopamine release induced by water intake during dehydration^37,42,44^. Importantly, however, inhibition of excitatory neurons in the SFO does not elicit dopaminergic neuron activity or dopamine release in the NAc^42^. Likewise, activation of inhibitory water-satiation neurons expressing GLP1 receptors in the SFO does not directly affect dopamine activity ^44^. These previous findings suggest that while activation of the thirst-driving system is a prerequisite for water intake to be rewarding and engage dopaminergic neurons, a mere reduction in the activity in this thirst system may not be sufficient to trigger increased activity in midbrain dopamine neurons. It has been reported that dopaminergic activation induced by water absorption from the gastrointestinal tract during dehydration requires the activity of inhibitory neurons in the lateral hypothalamus^44^. However, it is unclear whether this mechanism could also account for the rapid dopaminergic response that occurs within seconds during spontaneous drinking behavior. Another study suggests that neurons in various brain regions projecting directly to VTA dopaminergic neurons respond to reward in the same way as dopaminergic neurons^45^, suggesting that the rewarding effect of water may be induced through the activity of distributed brain circuits rather than a single brain region.

The acute rewarding effect of salt depends primarily on the peripheral tongue, and particularly requires an influx of sodium ions into specific taste receptor cells (TRCs) through the epithelial sodium channel (ENaC)^12,25^. In accordance with this, we showed that hedonic responses to salt reward in salt-depleted mice disappeared after amiloride treatment, an antagonist of ENaCs. Although the peripheral mechanism has been revealed, the neural circuitry responsible for its rewarding effect largely remains unknown. Salt craving has been demonstrated to be initiated by the activation of specific cells within distinct brain regions. Notably, the SFO, the pre-locus coeruleus, and the PBN have been found to play significant roles in salt craving ^6,11,12,46^. Furthermore, previous studies have revealed that input from the SFO to the bed nucleus of the stria terminalis (BNST) promotes salt craving, while input from the PBN to the BNST inhibits salt craving by activating inhibitory neurons in the BNST ^6,46^. The activity of specific projections from the BSNT to the VTA has been reported to control both reward and aversive behavior^47^, likely through the stimulation of distinct neural populations in the striatum^48,49^. Taken together, these studies suggest that the BNST may play a role in translating salt appetite into approach behavior.

In the present study, high salt intake in the dehydrated state and water intake in the desalted state each functioned as a punishment, suppressing consummatory licking and dopamine neuron activity. Previous research indicates that the perception of water is mediated by the activity of specific TRCs, suggesting that these TRCs play important roles in the aversive effect of excessive water ingestion^41^. In contrast, the aversive effects of high salt intake may depend on bitter and acid receptors^50^ as well as chloride channels^51^ rather than ENaCs in TRCs. Regarding the central mechanism, there exist several specific neuronal populations expressing specific genes such as calcitonin related polypeptide alpha (Calca), oxtocin receptor, and serotonin 2c receptor, within the PBN that exert an inhibitory effect on water- and salt-seeking behaviors^13,52,53^. Therefore, it is plausible that these cell populations in the PBN are important for the aversion that arises from excessive salt and water intake. Furthermore, based on previous findings that neural input from the PBN to the VTA decreases feeding behavior^54,55^, it is possible that the reduction in VTA dopamine neuron activity and dopamine levels in the striatum induced by excessive salt or water intake may be mediated by the PBN-VTA pathway.

Prior research has developed the homeostatic reinforcement learning (HRL) model to explain state-dependent learning and reward seeking behaviors^32^. Importantly, our previous study showed that this HRL model could explain salt-seeking behavior as well^31^. Furthermore, another study revealed that the HRL model can also explain enhanced dopamine release during salt intake under salt-deprived conditions by assuming a one-dimensional state space representing the internal sodium level^38^. However, it remains unclear whether the HRL can fully explain the state-dependent and flexible neural responses that either activate or suppress dopamine neurons. Indeed, a previous simulation study excluded testing dopaminergic responses to stimuli that exacerbate physiological needs, deeming such conditions as unsimulatable^38^. In the present study, we assumed a two-dimensional state space of water and salt in the HRL model to tackle this question. As a result, we found that (1) state-dependent behavioral preference and aversion to water or salt, (2) enhanced excitatory dopaminergic responses during water intake under water-deprivation and salt intake under salt-deprivation, and (3) that the suppression of dopaminergic activity during salt intake under water-deprivation and water intake under salt-deprivation, can all be qualitatively well-simulated by our HRL model. The results of our study suggest that the dopaminergic response to food intake is strongly dependent on homeostatic needs.

In summary, the current study revealed bidirectional excitatory and inhibitory responses of dopamine neurons in the VTA and dopamine release in the NAc during food intake depending on the internal salt-water balance in mice. Our results suggest that dopamine neurons control flexible approach and avoidance behaviors, as well as their learning, by encoding the value of food in a nutritional-state-dependent manner. These findings contribute to the elucidation of neural circuits controlling appetitive behaviors and their computational mechanisms, as well as the deeper understanding of the pathophysiology underlying psychiatric disorders associated with dysregulated appetite.

## METHODS

### Subjects

Sixteen DAT-cre mice (Slc6a3, The Jackson Laboratory; 9 males and 7 females) that were 8–12 weeks old and on a C57BL/6 JJcl background, and fourteen 8-week-old male C57BL/6JJcl mice (CLEA, Tokyo, Japan) were used. Mice weighed 20-25g and were provided food and water *ad libitum* until behavioral experiments were started. Mice were singly housed on a 12-h light/dark cycle (0800/2000), and experiments were carried out during the light cycle. All animal experiments were approved by the Animal Experimental Committee of the Institute for Protein Research, University of Osaka (29-02-1 and R04-01-2).

### Stereotaxic cannula implantation and virus injection

All surgeries were conducted under isoflurane gas anesthesia (1.5-3%) and using a stereotaxic frame (RWD Life Science, Shenzhen, China). For recording of dopamine neuron activity, 400 nl of AAV9-hSyn-FLEX-jGCaMP8s (2.7 × 10^13GC/ml, Addgene, #162377, MA, USA) mixed with 100 nl of AAV9-CAG-FLEX-tdTomato (3.8 × 10^13GC/ml, Addgene, #28306, MA, USA) was unilaterally injected into the VTA (coordinates in mm: AP – 3.3, ML ± 0.4 from bregma, and DV− 4.5 from the brain surface). For recording of dopamine release, 400 nl of AAV9-hSyn-GRABDA2m (2.5 × 10^13GC/ml, Addgene, #140553, MA, USA; 2.5 × 10^13GC/ml, WZ Bioscience Inc, MD, USA) mixed with 100 nl of AAV5-hSyn-mCherry (2.3 × 10^13 GC/ml, Addgene, #114472, MA, USA) was unilaterally injected into the nucleus accumbens core (AP + 1.4, ML ± 1.1 from bregma, and DV− 3.4 from the brain surface). In both experiments, injections were performed using a Nanoliter 2020 Injector (WPI, FL, USA) at a rate of 60 nl/min, and the injection glass capillary remained in place for 5–10 min to reduce backflow. Immediately after viral infusions, a 200 µm diameter (NA 0.37) optic fiber (RWD Life Science, Shenzhen, China) was implanted into the VTA (coordinates in mm: AP – 3.3, ML ± 0.4 from bregma, and DV− 4.3 from the brain surface) or nucleus accumbens core (AP + 1.4, ML ± 1.1 from bregma, and DV− 3.2 from the brain surface), and fixed in place with Superbond (Sunmedical, Shiga, Japan) and the dental cement (UNIFAST; GC, Tokyo, Japan).

### Behavioral experiments

#### Apparatus

Water- or salt-ingestive behavior was assessed in a metal operant chamber with a grid-floor and a drinking spout protruding from one sidewall (Med Associates, Inc., VT, USA). The chamber was illuminated by a house light. Our custom-made drinking spout had three holes through which three different liquid stimuli could be delivered. Liquid delivery through the spout was controlled by pinch valves (#HYN-3, CKD, Aichi, Japan). Licking responses to the feeding spout were counted with a contact lickometer (Med Associates, Inc., VT, USA). MED-PC software (Med Associates, Inc., VT, USA) was used for the quantification of spout-licking behavior and the control of valve opening for liquid delivery.

#### Test of sucrose and quinine

Water restriction started at least two days before pre-training and continued throughout the pre-training and test phases. Animals were allowed limited daily access to the water for 10 min in the home cage after daily pre-training or test sessions. During the pre-training, mice were trained on fixed ratio (FR) 10 schedule for 3 days where they had to lick the drinking spout 10 times to earn one drop of deionized water reward (10ul). The water rewards were delivered randomly through one of three holes on the spout. This 3-day FR10 training was followed by variable interval schedule (VI) training in which the water reward was delivered through the drinking spout as soon as mice exhibited spout-licking behavior after the inter-trial-interval (ITI) had passed. Each ITI was randomized with a uniform distribution between 10 s and 30 s, with a mean of 20 s (i.e. VI20 schedule). FR10 and VI20 trainings were finished when mice consumed 30 or 60 rewards, respectively, or after 30 min had passed. Once animals showed frequent licking for the water reward in the VI20 training, a single-drop brief access test was conducted. In this test, one of two different tastants, sucrose (300mM) or quinine (1mM) solution, was randomly delivered through different holes in the feeding spout. This test was conducted under a VI20 schedule and finished when mice consumed each tastant 20 times or after 60-min had passed.

#### Test of water and salt: water-restriction phase 1

Animals were water-restricted and trained in the operant-licking task to get deionized water for several days in the pre-training and then finally tested for taste reactivity to deionized water, 300mM, or 750mM NaCl solution delivered through 3 different holes in the drinking spout. Procedures for the water-restriction, pre-training, and testing were the same as those of the aforementioned test of sucrose and quinine.

#### Test of water and salt: salt-restriction phase

After finishing water-restriction phase 1, food and water in the home cage were replaced with a sodium-free diet (Oriental Yeast Co., LTD, Tokyo, Japan) and deionized water (Milli-Q water), respectively. One day before pre-training in the salt-restriction phase, mice were injected with furosemide (Nichi-iko, Toyama, Japan; 50mg/kg, i.p.) to facilitate the excretion of sodium from their body. In pre-training, mice were trained in the operant-licking task under a VI20 schedule for at least 3 days to earn a 300mM NaCl liquid. Once animals showed frequent operant-licking behavior, the test was conducted. Similar to the test under water-restriction, licking responses to Milli-Q water, 300mM, or 750mM NaCl solution were tested under salt-restriction.

#### Test of water and salt: water-restriction phase 2

After finishing the salt-restriction phase, animals were subjected to water-restriction again. In this phase, animals were trained in the operant-licking task under a VI20 schedule to get deionized water for several days, and then finally tested in the single-drop brief access test. Procedures of the water-restriction and testing were the same as those of water-restriction phase 1.

#### Behavioral measurement

During testing, each individual lick was measured. For the analysis of taste-evoked licking responses, we drew peri-event-time histograms (PETH) of the averaged licking frequency and calculated the area-under-the-curve (AUC). Furthermore, we also analyzed the microstructure of consummatory licks. A lick bout was defined as the first unit consisting of at least three consecutive licks and separated by a pause of 250ms after the reward delivery. We analyzed bout duration, licks per bout, and lick rate in bout^33,34^.

#### The effect of ENaC blockade

The epithelial sodium channel (ENaC) was blocked by mixing its antagonist, amiloride hydrochloride (Tokyo Chemical Industry Co., Ltd., Tokyo, JAPAN; 0.1mM), into each liquid delivered (i.e. water, NaCl 300mM and 750mM). In this test, mice were pretrained and tested under water-restriction and salt-restriction with or without amiloride.

#### Fiber photometry

Fluorescent signals were measured using a CMOS camera-based fiber photometry system (Doric Lenses Inc., QC, Canada) as described previously^56^. Briefly, fluorescence signals were obtained by exciting cells expressing jGCaMP8s or GRAB-DA2m with a 470 nm LED (30 µW at fiber tip), while dopamine-independent signals were obtained by exciting cells expressing tdTomato or mCherry with a 560 nm LED (30 µW at fiber tip). 470 nm and 560 nm LED lights were alternately pulsed at 12 Hz, and emitted lights were recorded using a CMOS camera, which acquired video frames of fluorescence from the end of the patch cord (NA = 0.37, 200 mm core; RWD Life Science, Shenzhen, China) at the same frequency. The timestamps between the fiber photometry and the behavioral chamber were synchronized using a TTL pulse which was generated from the behavioral chamber and sent to the fiber photometry system.

Video frames (fluorescent signals) were analyzed using Doric neuroscience studio ver. 6.2.3.0. (Doric Lenses Inc., QC, Canada). 470 nm-derived and 560 nm-derived signals were used for analysis. To compute the dF/F for signals, the raw signal was first baseline-corrected by subtracting its moving median, calculated using a 60-second sliding window. The resulting difference was then normalized by the same moving median, yielding dF/F = (raw signal – moving median) / moving median. This dF/F was z-scored by using the zscore function in Matlab and used for further analysis. For the analysis of tastant-evoked signal change, we drew PETHs of the averaged z-scored signal among animals, and also calculated positive and negative components of the AUC from 0 to 8 s after the onset of the first consummatory licking. Positive and negative AUCs were defined as the AUC above or below zero, respectively.

#### Histological verification

Animals were deeply anesthetized with ketamine (100mg/ kg) and xylazine (20mg/kg), and transcardially perfused with 0.01M PBS followed by 4% paraformaldehyde (PFA) in 0.1M PB (pH 7.4). Brains were removed and post-fixed with 4% PFA at 4 °C for 2 days. After cryoprotection, brains were embedded in O.C.T compound (Sakura Finetek Japan CO., Ltd, Tokyo, Japan) and cryosectioned at 40 µm. Sections were mounted with antifade mounting medium with DAPI (#ab104139, ABCAM, Cambridge, England). Images were acquired using a Keyence BZ-X800 microscope (Keyence, Osaka, Japan).

#### Statistics

All averaged data was expressed as the mean ± SEM. Raster plots of licking, heat maps of dopamine change, and PETHs of reward-related licking and dopamine change were created using Matlab software (MathWorks, CA, USA). All statistical analyses were performed using Prism (Graphpad, CA, USA) software. Normality of data distribution was checked by the Shapiro-Wilk test. Statistical methods are shown in each figure legend for each experiment. All ANOVAs were followed by post-hoc Tukey’s multiple comparisons test. For the analysis of the correlation between the number of licks and the dopaminergic response, Pearson’s correlation coefficient and liner regression analysis were performed. The F-test was used to assess whether the slope of the regression line was significantly different from zero. Scatter among replicates (i.e. trials) were taken into account. For all statistical tests, significance was assessed using an α value criteria of 0.05.

#### Computational modeling

To elucidate how variations in dopaminergic neuronal activity, which depends on variations in internal states of hydration and sodium levels, influence reward prediction and choice behavior, we employed a decision-making computational model. Specifically, we utilized the Homeostatic Reinforcement Learning (HRL, HRRL) model, a robust computational framework for representing internal state-dependent decision-making processes such as those influenced by hydration and sodium levels. Previous research has demonstrated that the HRL model could effectively explain how organisms maintain internal states such as adaptive levels of sodium and water through decision-making^31,32^. By simulating the dopamine changes and behavior of mice during the operant-licking task using the HRL model, we aimed to interpret the computational processes underlying changes in dopaminergic neuronal activity, which are greatly influenced by differences in the internal state, and to provide a functional explanation for these dynamics.

#### Computational model:Homeostatic Reinforcement Learning (HRL) model

The HRL model employed in this study is based on the assumption that homeostasis maintenance can be understood as a reinforcement learning process. It frames the minimization of deviations of internal states from their optimal levels as the calculation of action values that maximize the total reward. In the HRL model, a multidimensional metric space, where each dimension represents an internal state (e.g., hydration level or sodium concentration), is defined as the “homeostatic space” (a schematic image of the structure of the homeostasis space is shown in Fig.4b). Within this homeostatic space, the drive function D(H_t) is defined as the distance between the internal state of the i-th component at time t (e.g., hydration level or sodium concentration), H^i_t, and the ideal internal state H^*_i:

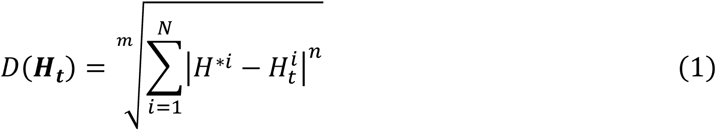

Here, m and n are free parameters that define the distance. As the internal state approaches the ideal state, the value of the drive function decreases. Based on this drive function, the reward r_t is determined as the change in the value of the drive function from time t to t+1. For example, if H_t represents the internal state of sodium, and K_t is defined as the amount of sodium ingested at time t, the relationship between the drive function and reward, derived from the internal state H_{t+1} at time t+1, is expressed as follows:

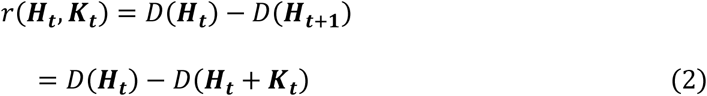

Furthermore, the natural decline in sodium balance is implemented using a time decay constant tau, as expressed in the following equation. The value of tau can be adjusted based on the internal state. When the internal states are set to represent water and sodium, the model is formulated as follows:

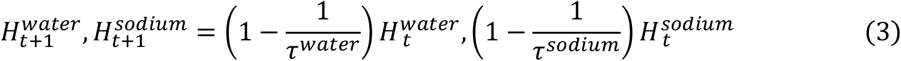

Combining these elements, the reward is calculated using the following equation:

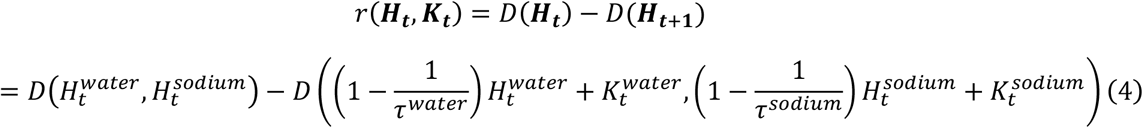

The update of action values based on reward prediction errors was modeled as a RL process using the Q-learning framework. In this model, the value Q_t(a) of an action a_t (e.g., water intake or taking no action) (Fig.4a) is updated based on temporal difference (TD) errors through the Q-learning ^31,32,57^ (Equation 5):

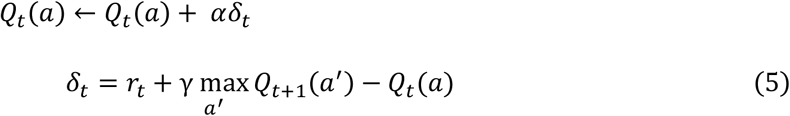

Here, alpha represents the learning rate for Q_t(a). Action selection depends on the relative magnitudes of the values (Q-values) for each action and follows a soft-max function:

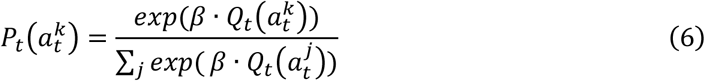

To facilitate a straightforward understanding of the HRL model algorithm, Fig.S9 illustrates a simulation of homeostasis maintenance for a one-dimensional internal state (e.g. water).

#### Simulation conditions and data Recording

Simulations were conducted under six experimental conditions, defined by the initial internal state (water- or salt-deprived) and the solutions used during training and testing. Specifically, mice began in a water-deprived state (WD-W) and were trained to lick for water, followed by a test in which only water was available. Under the next two water-deprived conditions, mice were similarly trained with water but tested either with 300 mM NaCl (WD-300) or with 750 mM NaCl (WD-750). The remaining three conditions involved mice that were initially salt-deprived and trained with 300 mM NaCl. These mice were then tested with either water (SD-W), 300 mM NaCl (SD-300), or 750 mM NaCl (SD-750).

For all six conditions, we recorded the time series data on internal states (H), behaviors (a), rewards (R), and reward prediction errors (TD error).

#### Training, Testing, and Measurement Procedures

In the three water-deprived conditions (WD-W, 300, 750), simulations began from a water-deprived state (H_0 = 100) and followed the same training protocol, in which subjects were trained to lick for water. Similarly, in the three salt-deprived conditions (SD-W, 300, 750), simulations started from a salt-deprived state (H_0 = 100), employed the same training regimen across all three conditions, this time training subjects to lick a 300 mM NaCl.

Each condition was followed by two tests, with only one solution offered during each test. We evaluated Q-values, the probability of intake behavior, actual behavior (a), intake volume (K), internal state (H), drive (D), reward (R), temporal difference errors (δ), and lick frequency. TD errors were quantified by pooling data from the two tests for each condition, as shown in Fig.3c. Licking behavior in the model was measured as the total number of “intake” behaviors in a single training or test trial (Fig. 4d) or the number of intake behaviors in a certain period of time (Fig. 4e-f), regardless of whether the liquid was acquired. Detailed simulation parameters are available in Supplementary Table 2.

## Supporting information

Supplementaly statistical data

## Data Availability Statement

The codes used in this study can be found at: https://github.com/YuukiUchida/WaterSaltDopamine_hRL

## ACKNOWLEDGEMENTS

This study was supported by the Japan Society for the Promotion of Science KAKENHI grants (JP21K15210 to T.M.; JP22H01105 to T.O.; JP20H00625 to Y.Y; JP23K24205, JP23K18163, and JP25K02547 to T.H.), Japan Agency for Medical Research and Development Grants (JP22gm6510012 and JP24wm0625111 to T.O.; JP21wm0425010 and JP21gm1510006 to T.H.), JST CREST (JPMJCR21P4 to Y.Y.), JST SPRING (JPMJSP2138 to IN, JPMJSP2120 to Y.U.), Salt Science Research Foundation Grants (2341 to T.O.; 2438 to T.H.), HOKUTO Foundation for the Promotion of Biological Science (to T.O.), LOTTE Foundation (to T.O.), Takeda Science Foundation (to T.O. and TH), Inamori Foundation (to T.O.), SR Foundation (to T.O.). Intramural Research Grant for Neurological and Psychiatric Disorders of NCNP (4-6, 6-9 to Y.Y.), the Collaborative Research Program of Institute for Protein Research, the University of Osaka (CRa-25-03) (to TH). The funders played no role in study design, data collection, analysis and interpretation of data, or the writing of this manuscript. Illustrations of animals in Fig.1d, 2c, 3b were created in BioRender.

## AUTHOR CONTRIBUTIONS

These authors contributed equally: T.O., I.N., Y.U.; Conceptualization: T.O, Y.Y, T.H; Data curation: T.O., I.N., Y.U.; Formal analysis: T.O., I.N., Y.U., M.A.; Funding acquisition: T.O., I.N., Y.U., T.M., Y.Y., T.H.; Investigation: T.O., I.N., Y.U.; Methodology: T.O., Y.U., Y.Y., T.H.; Project administration: T.O., Y.Y., T.H.; Supervision: T.O., Y.Y., T.H.; Writing - original draft: T.O., I.N., Y.U., T.M., Y.Y., T.H.; Writing-review and editing: T.O., I.N., Y.U., T.M., Y.Y., T.H.; All authors read and approved the final manuscript.

## ADDITIONAL INFORMATION

Supplementary information: The online version contains supplementary material.

