## Supplementaly information for "Flexible value coding in the mesolimbic dopamine system depending on internal water and sodium balance"

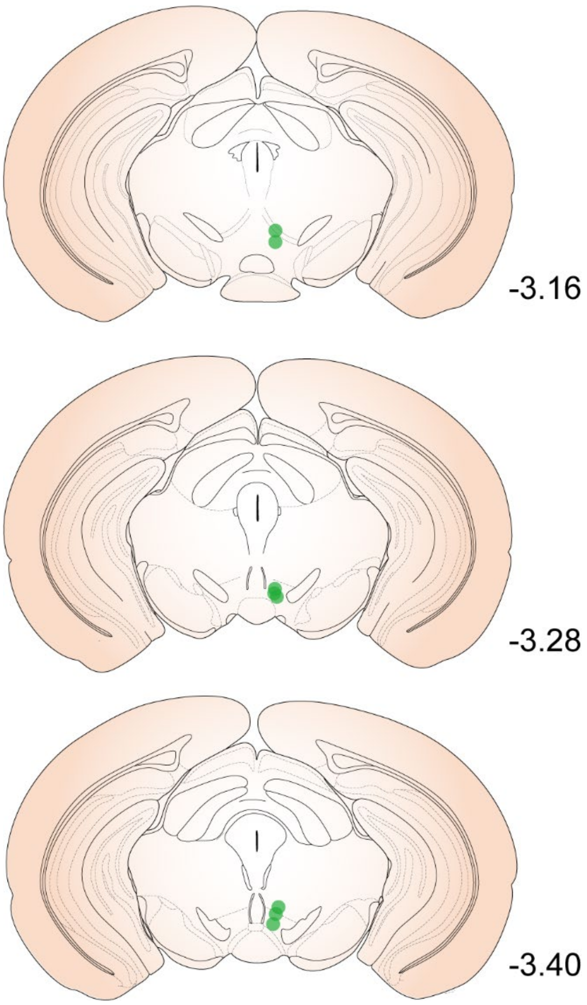


Supplementary Figure S1. Optical fiber placements.

Histology of optical fiber placements for the imaging of dopamine neurons in the VTA in the experiment for sweetness- and bitterness-evoked response. Green circles show tip placement of optical fibers in each mouse (n=8).

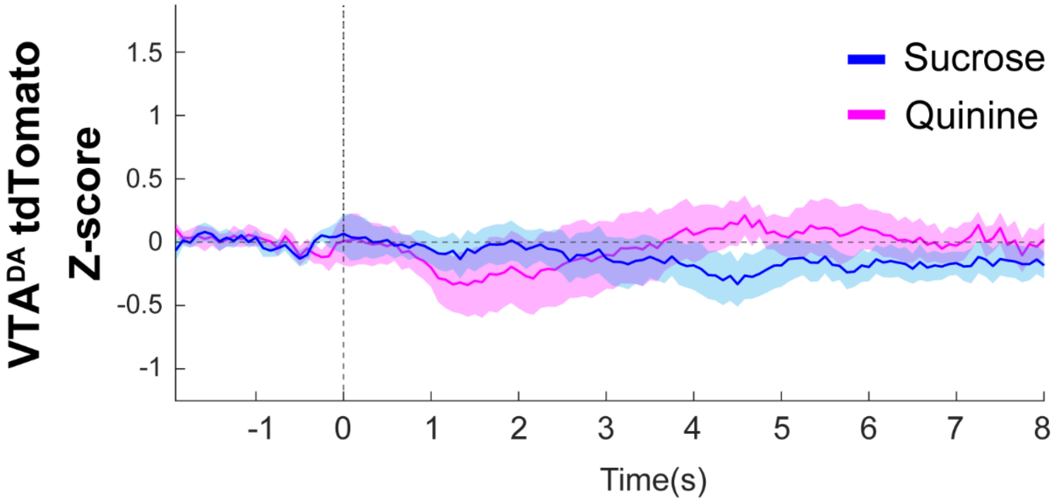


Supplementary Figure S2. Signal of red fluorescent proteins as a control.

PETHs of the tdTomato signal in dopamine neurons in the VTA.

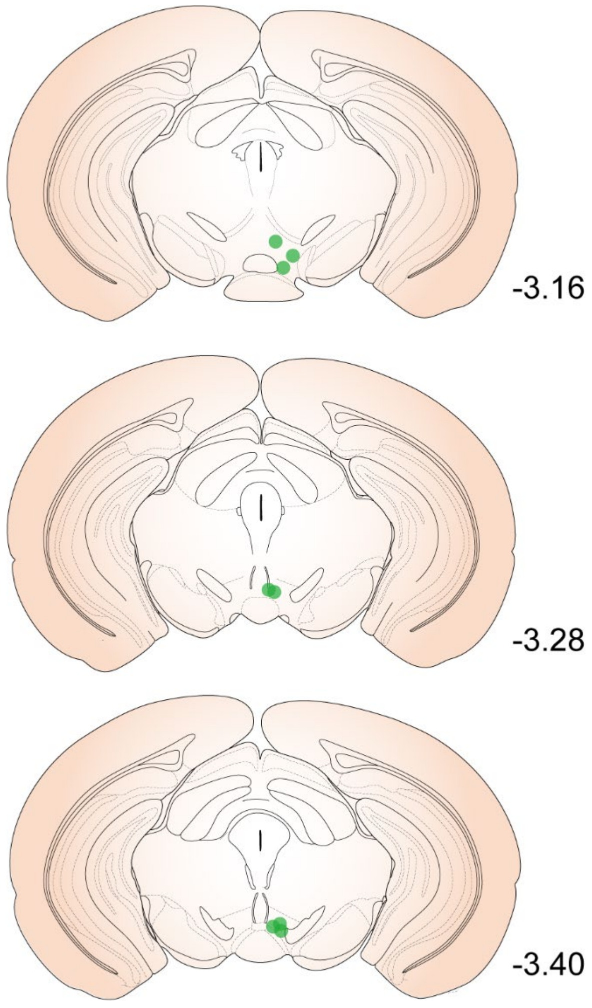


Supplementary Figure S3. Optical fiber placements.

Histology of optical fiber placements for the imaging of neural responses of dopamine neurons in the VTA to water and salt intake. Green circles show the tip placement of optical fibers in each mouse (n=8).

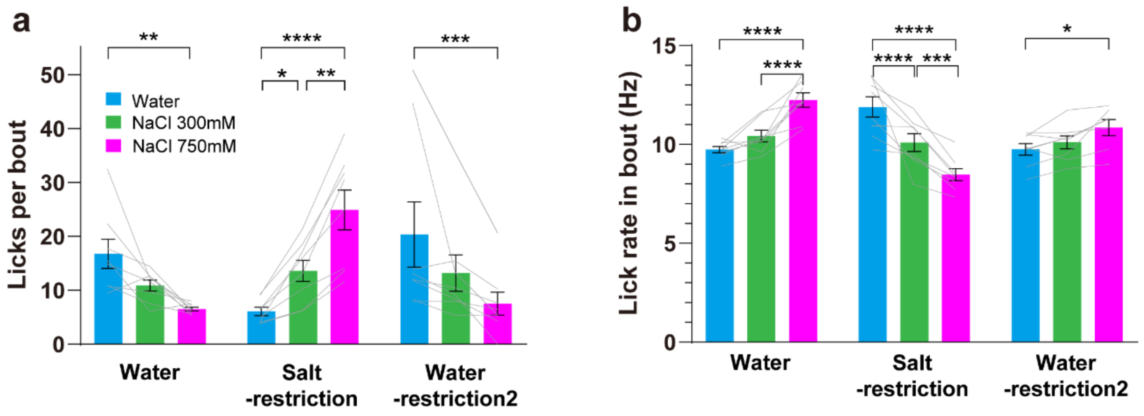


Supplementary Figure S4. State-dependent changes in licking responses to water and salt during recording of VTA dopamine neuron activity.

a Licks per bout. b Lick rate in bout (Hz). Gray lines overlayed on bar plots indicate data from individual animals. a,b post-hoc Tukey's multiple comparisons test. **** p<.0001, *** p<.001, ** p<.01, and *p<.05.

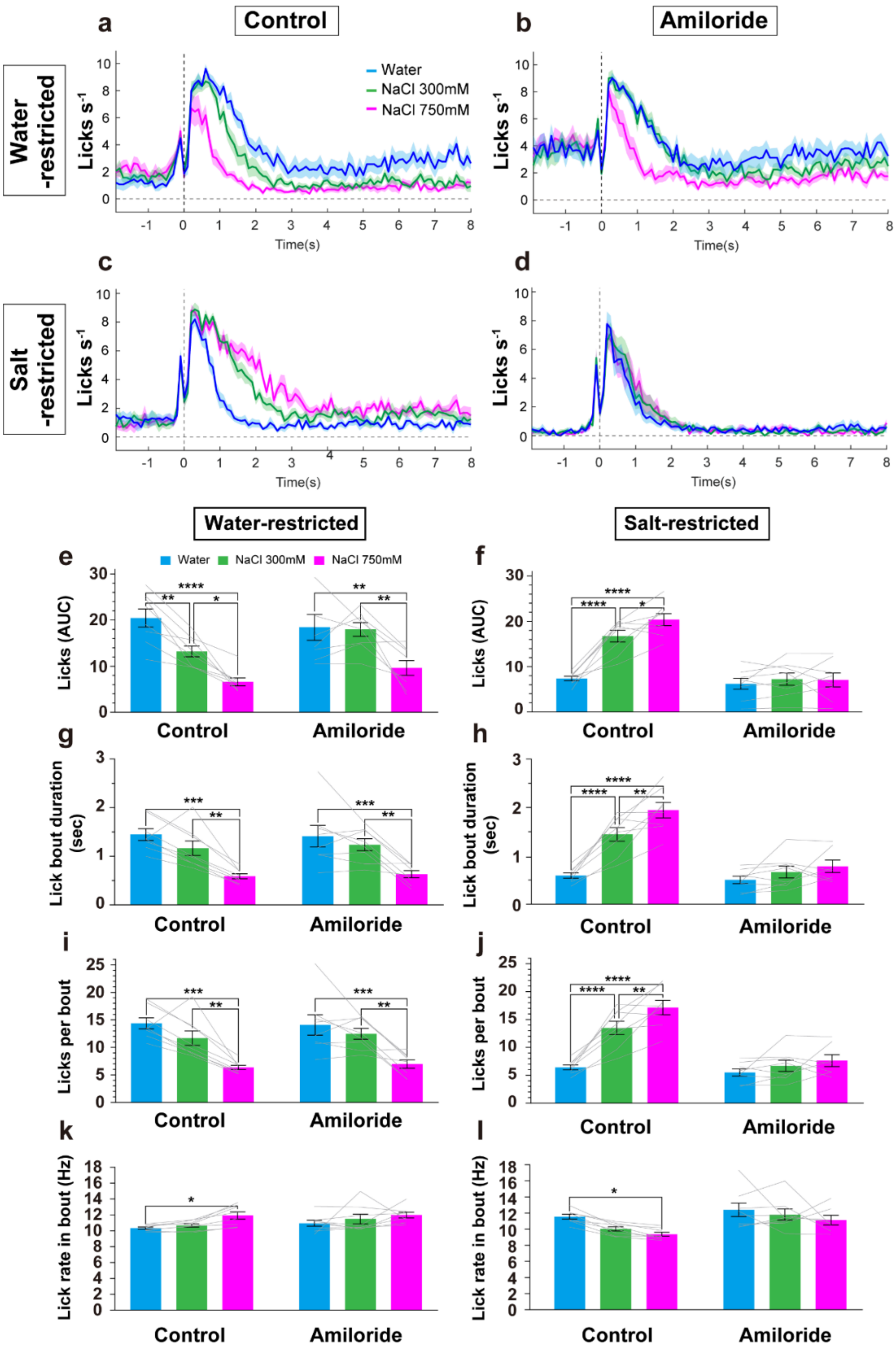


Supplementary Figure S5. The effect of amiloride on water and salt intake.

Amiloride greatly reduced rewarding salt intake, but the effect on salt-aversion was limited. a-d PETH of licking responses with or without amiloride under water- or salt-restriction. e, f Licks (AUC) under water-restriction (e) or salt-restriction (f), respectively. g, h Lick bout duration (sec) under water-restriction (g) or salt-restriction (h), respectively. i, j. Licks per bout under water-restriction (i) or salt-restriction (j), respectively. k, l Lick rate in bout under water-restriction (k) or salt-restriction (l), respectively. Gray lines overlayed on bar plots indicate data from individual animals. e, f, g, h, I, j, k, l, post-hoc Tukey's multiple comparisons test. N=8. **** $p<.0001$, *** $p<.001$, ** $p<.01$, and *$p<.05$.

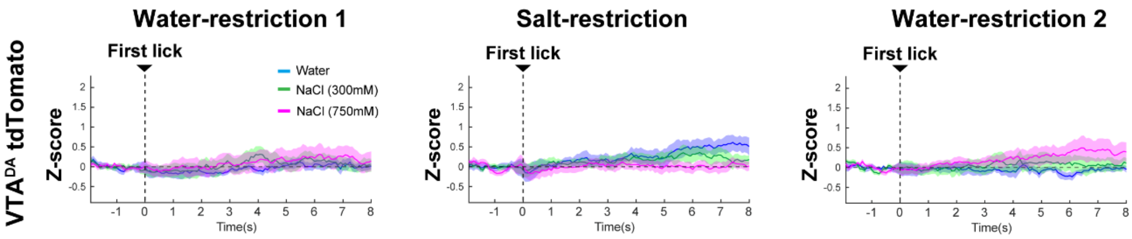


Supplementary Figure S6. Signal of red fluorescent proteins as a control.

PETHs of the tdTomamo signal from dopamine neurons in the VTA under water-restriction 1 (left), salt-restriction (middle), and water-restriction 2 (right).

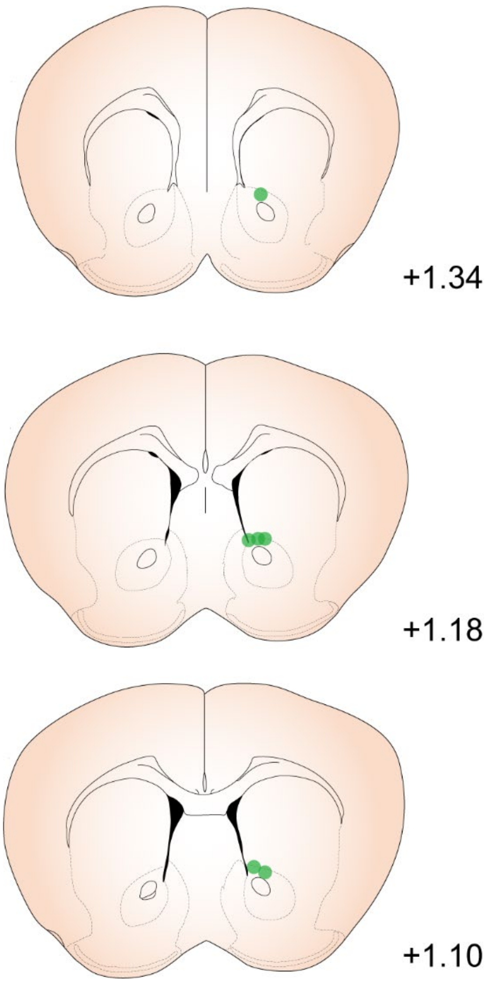


Supplementary Figure S7. Optical fiber placements.

Histology of optical fiber placements for the imaging of dopaminergic responses in the NAc to water and salt intake. Green circles show the tip placement of optical fibers in each mouse (n=6).

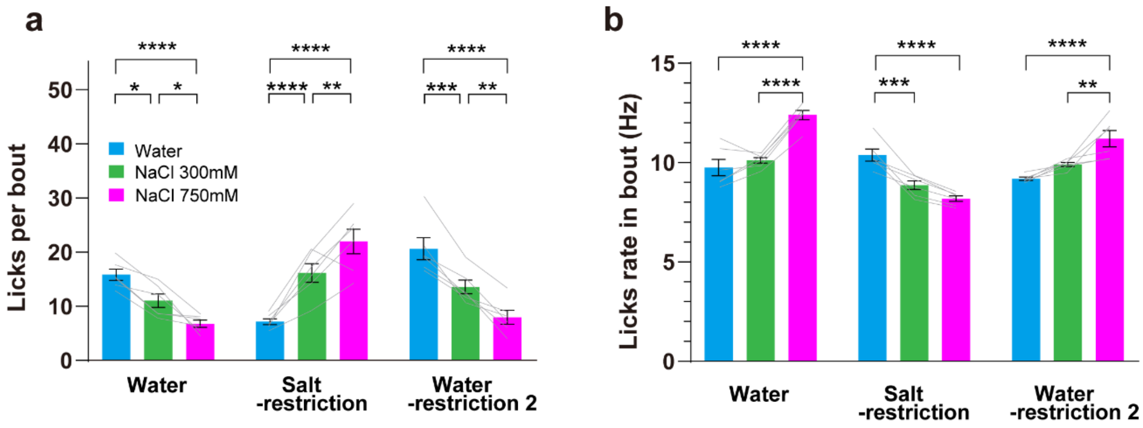


Supplementary Figure S8. State-dependent changes in licking responses to water and salt during recording of dopamine dynamics in the NAc.

a Licks per bout. b Lick rate in bout (Hz). Gray lines overlayed on bar plots indicate data from individual animals. a,b post-hoc Tukey's multiple comparisons test. **** p<.0001, *** p<.001, ** p<.01, and *p<.05.

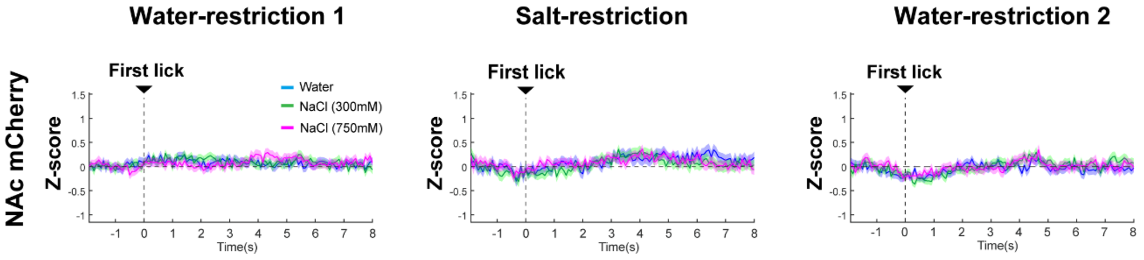


Supplementary Figure S9. Signal of red fluorescent proteins as a control.

PETHs of the mCherry signal from the NAc under water-restriction 1 (left), salt-restriction (middle), and water-restriction 2 (right).

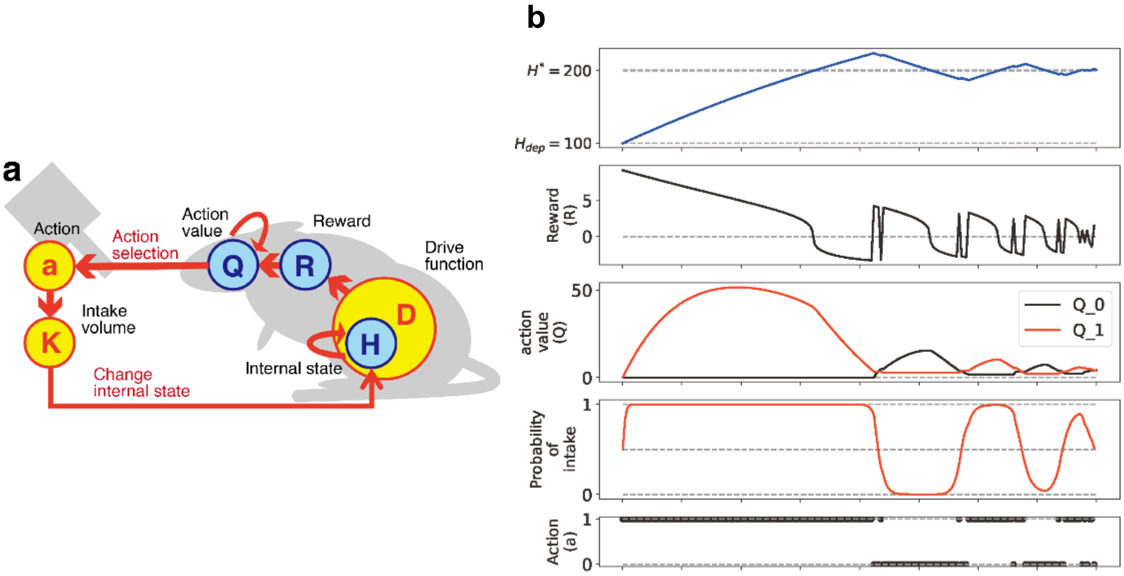


Supplementary Figure S12. Simplified algorithm of the one-dimensional HRL model.

a, b A schematic representation of the algorithm is shown. An example of time-series data from a learning simulation is presented, in which a one-dimensional internal state (e.g., water) starts in a deprived initial state ($H_0 = 100$, dotted line) and continuous action selection occurs between "do nothing" and "water intake." This simulation assumes only a one-dimensional internal state. The external state ($S_0$) and two actions, "do nothing" ($a_0$) and "water intake" ($a_1$), were evaluated (Fig. 4a). Action selection depends on the relative magnitudes of the values (Q-values) for each action and follows a soft-max function (see equation 6). In the supplementary simulation, the initial Q-values for both actions were set to 0 (Q_t = [0, 0]). Therefore, the first action was selected randomly. When the intake action was executed, the internal sodium state increased according to the constant parameter $K_t$, which defines the amount of water intake. If "do nothing" was selected, $K_t$ was set to zero ($K_t = 0$). At t = 0, the initial internal state ($H_0 = 100$) was set significantly lower than the ideal state ($H^* = 200$, dotted line) to represent water deprivation. At this stage, the drive function value was high, as it represents the distance between the internal state ($H_t$) at time t and the ideal state ($H^* = 200$) (see equation 1). When the agent performed the intake action, the internal state increased, reducing the drive function value and resulting in a positive reward (see equation 2). To update the *Q*-values based on reward values, *Q*-learning was used. In this model, *i* represents each action, alpha is the learning rate, and $\delta_t = R_t + \gamma \max_{a'} (Q_{t+1}(a')) -$

$Q_t(a)$ represents the TD error. After updating the Q-values, the agent selected the next action. Following random action selection, the Q-value for intake increased, and the internal nutritional state rapidly reached the ideal point, maintaining behavioral homeostatic regulation. The value of "do nothing" decreased, and the Q-value for intake remained relatively high even after exceeding the set value ($H_t > H^* = 200$). Subsequently, the value of "do nothing" exceeded the Q-value for intake due to the excess internal state, leading to an increased frequency of "do nothing". Continued "do nothing" behavior caused a natural decay in the internal state. In response to this decline, the Q-value for intake exceeded the Q-value for "do nothing", resulting in sustained homeostatic regulation. Detailed parameter values for the simulation are provided in Table S2. The lines relating to intake actions are depicted in red-orange.

Supplementary Table S1; Simulation parameters used in Figure 3

| Free parameter | Value | Explanation |
|---|---|---|
| $\alpha$ | 0.001 | Learning rate of state-action values |
| $\beta$ | 0.05 | Inverse temperature of action selections |
| $\gamma$ | 0.7 | Discount rate |
| $m$ | 1 | Free parameter of the drive function |
| $n$ | 2 | Free parameter of the drive function |
| $\tau_{water}$ | 7000 | Attenuation rate of the internal water state |
| $\tau_{sodium}$ | 5000 | Attenuation rate of the internal sodium state |
| $K_{water}$ | 22 | Volume of water in an intake |
| $K_{300mM\ NaCl}$ | 35 | Volume of sodium in a 300 mM intake |
| $K_{750mM\ NaCl}$ | 100 | Volume of sodium in a 750 mM intake |
| $H^*$ | 1000 | The ideal internal state |
| $H_{dep}$ | 700 | The depleted initial internal state |

Supplementary Table S2; Simulation parameters used in Figure S3

| Free parameter | Value | Explanation |
| --- | --- | --- |
| $\alpha$ | 0.3 | Learning rate of state-action values |
| $\beta$ | 0.6 | Inverse temperature of action selections |
| $\gamma$ | 0.9 | Discount rate |
| $m$ | 3 | Free parameter of the drive function |
| $n$ | 4 | Free parameter of the drive function |
| $\tau$ | 200 | Attenuation rate of the internal state |
| $K$ | 2 | Volume of small intake |
| $H^*$ | 200 | The ideal internal state |
| $H_0$ | 100 | The depleted initial internal state |
